# Repeated ethanol exposure and withdrawal alters ACE2 expression in discrete brain regions: Implications for SARS-CoV-2 infection

**DOI:** 10.1101/2022.03.29.486282

**Authors:** Nagalakshmi Balasubramanian, Thomas D James, Selvakumar Govindhasamy Pushpavathi, Catherine A. Marcinkiewcz

## Abstract

Emerging evidence suggests that people with alcohol use disorders are at higher risk for SARS-CoV-2. SARS-CoV-2 engages angiotensin-converting enzyme 2 (ACE2) and transmembrane serine protease 2 (TMPRSS2) receptors for cellular entry. While ACE2 and TMPRSS2 genes are upregulated in the cortex of alcohol-dependent individuals, information on expression in specific brain regions and neural populations implicated in SARS-CoV-2 neuroinvasion, particularly monoaminergic neurons, is limited. We sought to clarify how chronic alcohol exposure affects *ACE2* and *TMPRSS2* expression in monoaminergic brainstem circuits and other putative SARS-CoV-2 entry points. C57BL/6J mice were exposed to chronic intermittent ethanol (CIE) vapor for 4 weeks and brains were examined using immunofluorescence. We observed increased ACE2 levels in the olfactory bulb and hypothalamus following CIE, which are known to mediate SARS-CoV-2 neuroinvasion. Total ACE2 immunoreactivity was also elevated in the raphe magnus (RMG), raphe obscurus (ROB), and locus coeruleus (LC), while in the dorsal raphe nucleus (DRN), ROB, and LC we observed increased colocalization of ACE2 with monoaminergic neurons. ACE2 also increased in the periaqueductal gray (PAG) and decreased in the amygdala. Whereas ACE2 was detected in most brain regions, TMPRSS2 was only detected in the olfactory bulb and DRN but was not significantly altered after CIE. Our results suggest that previous alcohol exposure may increase the risk of SARS-CoV-2 neuroinvasion and render brain circuits involved in cardiovascular and respiratory function as well as emotional processing more vulnerable to infection, making adverse outcomes more likely. Additional studies are needed to define a direct link between alcohol use and COVID-19 infection.

## Introduction

The outbreak of coronavirus disease-2019 (COVID-19) caused by severe acute respiratory syndrome coronavirus 2 (SARS-CoV-2) has spread rapidly around the world and is a major global health concern. SARS-CoV-2 can infect the brain and cause neurological symptoms in patients that include headache, dizziness, anosmia, ageusia, ataxia, loss of autonomic respiratory control, lethargy, depression, and anxiety (Haidar et al., 2021; Kabbani and Olds, 2020; Mao et al., 2020; Nagu et al., 2021; Satarker and Nampoothiri, 2020). Importantly, the COVID-19 pandemic coincides with a rise in substance use disorders (SUD) (Alexander et al., 2020; Becker and Fiellin, 2019; Volkow, 2020), but whether previous exposure to drugs of abuse confers additional risk of SARS-CoV-2 infection or neurological complications is unknown. As SARS-CoV-2 may infect brain circuits that regulate respiratory control, the combination of a history of drug and alcohol use and COVID-19 infection could be particularly lethal (Wang et al., 2021). Chronic use of opioids, alcohol and other drugs is also associated with cardiovascular, pulmonary, metabolic (Bahorik et al., 2017; Friedman et al., 2006, 2003) and immune-related (Barr et al., 2016) diseases, all of which are potent risk factors for COVID-19. Consequently, it stands to reason that individuals with SUD, opioid use disorders (OUD), or alcohol use disorder (AUD) would be at heightened risk for adverse COVID-19 outcomes, including mortality. Currently, research is scarce on the effects of addictive drugs including cannabis, cocaine, and alcohol on the susceptibility to COVID-19 (Volkow, 2020).

It is well-established that angiotensin-converting enzyme-2 (ACE2), a significant player in the renin-angiotensin system (RAS), is the key receptor for SARS-CoV-2 invasion (Wan et al., 2020; Zhou et al., 2020). The virus binds to ACE2 via its trimeric spike (S) protein which is cleaved by the transmembrane serine protease 2 (TMPRSS2), triggering the fusion of viral and cellular membranes and internalizing the virus into the cell (Zhang et al., 2020). ACE2 is then shed from the membrane producing soluble ACE2 (sACE2) (Wang et al., 2022). Recent studies have confirmed that sACE2 can also act as a receptor for SARS-COV-2, mediating viral entry into the cell and then propagating to the infected area (Wang et al., 2022). Consequently, ACE2 deficiency after viral invasion might augment the dysregulation between the ‘adverse’ ACE→Angiotensin II→AT1 receptor axis and the ‘protective’ ACE2→Angiotensin 1-7→Mass receptor axis, leading to inflammation, vasoconstriction, hypertension, and fibrosis (Verdecchia et al., 2020).

Invasion and propagation of SARS-CoV-2 into the central nervous system (CNS) has been demonstrated explicitly in post-mortem tissues from COVID-19 patients and preclinical studies in transgenic mice expressing human *ACE2* (Song et al., 2021). Furthermore, neurodegeneration and brain edema were also observed in the autopsies of COVID-19 patients (Xu et al., 2020). Mounting evidence suggests that ACE2 is robustly expressed in nuclei involved in the central regulation of cardiovascular function like the cardio-respiratory neurons of the hypothalamus and brainstem, as well as in non-cardiovascular areas such as the motor cortex and raphe (Doobay et al., 2007; Hernández et al., 2021; Lin et al., 2008; Lukiw et al., 2022; Qiao et al., 2020). ACE2 is also prominent in astrocytes, pericytes, and endothelial cells (Gallagher et al., 2006; Hernández et al., 2021) representing its role in neuroinflammation after COVID-19 infections. Altogether, the evidence suggests that the pattern and density of *ACE2* expression can affect the risk of SARS-CoV-2 neuroinvasion and propagation within the brain, which in turn may lead to lethal cardiorespiratory complications as well as acute and long-term neurological sequelae.

Recent observations have shown that COVID-19 associated genes including *ACE2*, and *TMPRSS2* were upregulated in the motor and frontal cortex of chronic alcoholics (Muhammad et al., 2021). In addition, expression of *ACE2* was observed in brainstem regions that contain dopaminergic, noradrenergic, and serotonergic neurons in rats (Hernández et al., 2021). Chronic alcohol exposure is known to affect the function and output of monoaminergic neurons dependence (Jaatinen et al., 2013; Ketcherside et al., 2013; Lowery-Gionta et al., 2014; Marcinkiewcz et al., 2016a, 2014), which may alter *ACE2* and *TMPRSS2* expression in these neurons and increase their susceptibility to SARS-CoV-2 infection. Here we test the hypothesis that chronic intermittent ethanol exposure (CIE) alters *ACE2* and *TMPRSS2* expression in various brain stem nuclei (locus coeruleus, raphe nucleus, periaqueductal gray) as well as other interconnected regions such as the hypothalamus, thalamus, and amygdala that are implicated in stress and reward processing. We also looked at regions such as the olfactory bulb that are implicated in SARS-CoV-2 entry into the brain(Bilinska et al., 2020; Lima et al., 2020; Nampoothiri et al., 2020; Netland et al., 2008). Here we used CIE vapor inhalation to induce ethanol dependence and withdrawal (Marcinkiewcz et al., 2014; Pati et al., 2020) and measured ACE2 and TMPRSS2 levels in monoaminergic neurons and associated brain nuclei by immunofluorescence.

## Materials and methods

### Animals

All experiments were performed on adult male C57BL/6J mice (3-6 months) following the ethical guidelines of NIH guidelines for animal research and approved by the Institutional Animal Care and Use Committee (IACUC) at the University of Iowa. Animals were housed in a ventilated and temperature-controlled vivarium on a standard 12-hour cycle (lights on at 0230) with *ad libitum* access to food (Envigo NIH-31 Modified Open Formula 7913) and water.

### Chronic intermittent ethanol (CIE) exposure

Chronic intermittent ethanol exposure was achieved via vapor inhalation as previously described (Marcinkiewcz et al., 2014). Briefly, mice were placed in standard mouse cages in Plexiglas® vapor chambers (30 × 10 × 15 cm, La Jolla Alcohol Research Inc., La Jolla, CA, USA) and exposed to ethanol volatilized by heating 95% ethanol mixed with fresh air at a rate of ≈10 L/min. To facilitate intoxication and stabilize blood ethanol concentration (BECs), mice were injected daily with an alcohol dehydrogenase inhibitor, pyrazole (Sigma, St. Louis, MO, USA; dosage: 1 mmol/kg, i.p.) and a loading dose of EtOH (1.57 g/kg) immediately before placement in vapor chambers. Air controls received only pyrazole and were placed in an alternate cage in the same room and exposed to room air. Each cycle of CIE or air exposure lasted 16 h per day (in at 1700 h, out at 0900 h), followed by an 8 h withdrawal for 4 consecutive days of exposure followed by an 80-h withdrawal. This cycle was repeated for a total of 4 weeks and the mice were perfused to collect brains for further immunostaining procedures.

### Reverse Transcriptase-Quantitative PCR (RT-qPCR)

A separate group of mice was decapitated under an isoflurane chamber to collect brains for RT-qPCR experiments. The brains were extracted and the olfactory bulb (OB), orbitofrontal cortex (OFC), nucleus accumbens (NAc), amygdala (AMY), thalamus (TH), hypothalamus (HT), hippocampus (HP), dorsal raphe nucleus (DRN), locus coeruleus (LC), and nucleus of the solitary tract (NTS) were dissected. Total RNA from the above-mentioned brain regions was isolated using TRIzol reagent (Ambion, USA) followed by DNA-free™ DNA Removal Kit (Life Technologies, USA) as described previously (Balasubramanian et al., 2021). RNA was then quantified using NanoDrop 1000 Spectrophotometer (Thermo Scientific, USA). The reverse transcription was performed using an iScript cDNA synthesis kit (Bio-Rad Laboratories, CA, USA) using the thermal profile: 25 °C for 5 min, 45 °C for 20 min, and 95 °C for 1 min.

Quantitative reverse transcription PCR (RT-qPCR) for the target gene β*-actin*, *ACE2*, and *TMPRSS2* was performed using SYBR green qPCR master mix (Bio-Rad Laboratories, USA) on CFX96™ Real-time-PCR System (Bio-Rad Laboratories, USA). The specific primer sets used for amplification of target cDNA are listed in Table 1. The thermal profile used for RT-qPCR were as follows: 95 °C for 10 min followed by 40 cycles of 95 °C for 30 s, 60 °C for 30 s, followed by a melt curve analysis profile: 60 °C to 95 °C in 0.5 °C increments at a rate of 5 s/step. Relative mRNA levels of *ACE2* and *TMPRSS2* in different brain regions were compared to OB and the fold change in the mRNA levels was determined after normalization to *β-actin* using the 2^−ΔΔCT^ method (Schmittgen and Livak, 2008). Results are represented as fold changes in the mRNA levels (± SEM).

### Western Blot

A separate group of mice was decapitated under an isoflurane chamber to collect brains for western blot experiments as reported previously (Abooj et al., 2016; Sagarkar et al., 2021). Briefly, different brains regions were dissected (as listed above) including raphe magnus and obscurus, for the western blot of β-actin, ACE2, and TMPRSS2. Total protein from all the tissues was isolated using RIPA lysis buffer. The proteins were quantified using the BCA method (ThermoFisher Scientific, USA) and an equal quantity of proteins was resolved in a 10% SDS-PAGE gel. According to the standard procedures, resolved proteins were further transferred to a PVDF membrane (0.45 μm; Millipore, USA) for immunoblotting and the blots were blocked using Starting Block T20 (TBS) Blocking Buffer (ThermoFisher Scientific, USA) for 1 h at room temperature. Further, the membranes were incubated overnight with antibodies specific to ACE2, TMPRSS2, and β-actin (Table 2) at 4°C. Secondary antibodies (IR dye; Table 2) were used to detect the primary antibodies. The blots were visualized, and the images were acquired using Sapphire Biomolecular Imager (Azure Biosystems Inc, USA) at 700 nm and 800 nm infrared. Protein bands were quantified using Image J 1.45 software (National Institutes of Health, Bethesda, MD). The average relative density of the ACE2 and TMPRSS2 was determined after normalization to β-actin. Results are represented as a mean relative density of the protein levels (± SEM).

### Immunofluorescence

ACE2 expression was quantified across specific brain regions using standard immunofluorescence methods. Expression in serotonergic or dopaminergic/noradrenergic neurons was quantified using tryptophan hydroxylase 2 (TPH2) or tyrosine hydroxylase (TH) immunofluorescence, respectively, as markers for identification. Briefly, mice were anesthetized with Avertin and transcardially perfused with phosphate-buffered saline (PBS), followed by 4% paraformaldehyde (PFA). The brains were extracted, post-fixed in PFA overnight, transferred serially to sucrose solutions (10, 20, and 30%), and then cut using a cryomicrotome (Leica Microsystems, Wetzlar, Germany) to a thickness of 20 µm. For each region of interest (ROI), 4-5 sections were used across the rostral-caudal axis. Sections were washed in PBS and permeabilized in 0.5% Triton X-100/PBS for 30 min, followed by blocking in 10% normal donkey serum in 0.1% Triton X-100/PBS and then incubated with the respective primary and secondary antibodies (Table 3) according to standard immunostaining procedures. Images were acquired on an Olympus FV-3000 confocal microscope in 0.5 µm steps and converted to maximum projection images using Image J software. For every ROI, multiple tiles were imaged, and mosaic stitching was performed using FV3000 Multi-Area Time-Lapse (MATL) Software. Images were analyzed by trained researchers blind to experimental conditions for expression parameters in ImageJ (Fiji) software as described previously (Sagarkar et al., 2021). Immuno-colocalization was analyzed by measuring the percentage of co-localization area pixels by total immunoreactive area pixels after setting color thresholding parameters in ImageJ (Fiji) software (Sagarkar et al., 2021). All the results analyzed were represented as mean percentage immunoreactive area (±SEM). To measure the degree of colocalization between ACE2 and neuronal markers, the Pearson correlation coefficient (PCC) and Mander’s overlap coefficient (MOC) was calculated using the Just Another Colocalization plugin (JACoP) in ImageJ (Fiji) software for all the colocalization analyses (Dunn et al., 2011). The results for both the coefficients were represented as the mean of the correlation coefficient values (±SEM).

### Data and statistical analysis

The statistical analysis was performed using GraphPad Prism v.9 (CA, USA). The significance of differences between the two groups (e.g. Air control vs. CIE) was assessed using the Student’s Unpaired t-test with Welch’s correction. The analysis for differential *ACE2* and *TMPRSS2* mRNA expression relative to the olfactory bulb was performed using a one-way analysis of variance (ANOVA). The levels of non-cleaved cellular ACE2 (cACE2) and sACE2 across different brain regions relative to the olfactory bulb were analyzed using a two-way ANOVA and TMPRSS2 protein levels were analyzed using one-way ANOVA. Post hoc analysis for all the ANOVA comparisons was carried out using Dunnett’s or Sidak’s multiple comparisons test. A p-value less than 0.05 was considered significant for all the analyses.

## Results

### Initial Findings: Differential ACE2 and TMPRSS2 expression in the CNS

SARS-CoV-2 invasion in the CNS via the nasal cavity, followed by entry into olfactory bulb cells and subsequent spread to deeper brain regions has been shown previously (Burks et al., 2021; Jiao et al., 2021; Song et al., 2021; Ueha et al., 2020; Xydakis et al., 2021). We measured *ACE2* expression in a subset of brain regions that are involved in either viral invasion or whose functions are impacted by CIE. Therefore, we measured *ACE2* expression in the olfactory bulb, monoaminergic nuclei, or brain regions that receive substantial innervation from either locus. After isolating tissue punches containing a region of interest, we quantified *ACE2* or *TMPRSS2* mRNA levels by performing RT-qPCR (Figure 1A and B). Figure 1A shows heterogenous *ACE2* mRNA levels in the discrete brain regions in which we were able to detect expression (one-way ANOVA main effect: F_9,30_ = 12.37, *p* < 0.001). Compared to olfactory bulb transcript levels, *ACE2* expression was significantly lower in the OFC, NAc, HP, AMY, and DRN (Dunnett’s multiple comparisons test: t_30_ = 4.532, *p* = 0.0007 for OFC; t_30_ = 4.210, *p* = 0.0016 for NAc; t_30_ = 3.567, *p* = 0.0090 for HP; t_30_ = 3.438, *p* = 0.0125 for AMY; and t_30_ = 4.087, *p* = 0.0023 for DRN). However, TMPRSS2 expression was restricted to a subset of these ACE2-expressing regions (Figure 1B). *TMPRSS2* mRNA expression was also not uniform (one-way ANOVA main effect: F_4,15_ = 6.175, *p* = 0.0038), with mRNA levels significantly lower in TH compared to OB (Dunnett’s multiple comparisons test: t_15_ = 2.771, *p* = 0.0460). In parallel, we also tested for ACE2 and TMPRSS2 protein in these brain regions. We detected both cACE2 and sACE2 in each of the regions that showed robust mRNA expression. Analysis compared to OB revealed significant differences in protein levels across loci, but also differences in soluble/cellular ACE2 levels (two-way ANOVA, main effect: F_11,72_ = 3.667, *p* = 0.0004 for loci and F_1,72_ = 15.22, *p* = 0.0002). Relative protein expression across brain loci were not identical to mRNA levels, with sACE2 being lower in HT, AMY, DRN, RMG, ROB, LC, and NTS (Sidak’s multiple comparisons test: t_72_ = 3.332, *p* = 0.0149 for HT; t_72_ = 4.033, *p* = 0.0015 for AMY; t_72_ = 3.924, *p* = 0.0022 for DRN; t_72_ = 4.642, *p* = 0.0002 for RMG; t_72_ = 4.782, *p* < 0.0001 for ROB; t_72_ = 3.819, *p* = 0.0031 for LC; and t_72_ = 3.331, *p* = 0.0149 for NTS) as compared to OB. Across brain regions, no differences in cACE2 levels were detected. Surprisingly, we also detected TMPRSS2 protein, even in regions such as the OFC, NAc, HP, and AMY where mRNA levels were too low to detect. Relative TMPRSS2 protein expression across brain loci were significantly lower in NAc, HP, TH, HT, RMG, ROB, and NTS as compared to OB (one-way ANOVA main effect: F_11,24_ = 4.405, *p* = 0.0012; Dunnett’s multiple comparisons test: t_24_ = 3.723, *p* = 0.0089 for NAc; t_24_ = 3.091, *p* = 0.0382 for HP; t_24_ = 3.764, *p* = 0.0081 for TH; t_24_ = 3.132, *p* = 0.0349 for HT; t_24_ = 4.259, *p* = 0.0024 for RMG; t_24_ = 3.381, *p* = 0.0199 for ROB; and t_24_ = 3.265, *p* = 0.0259 for NTS).

**Figure 1:**
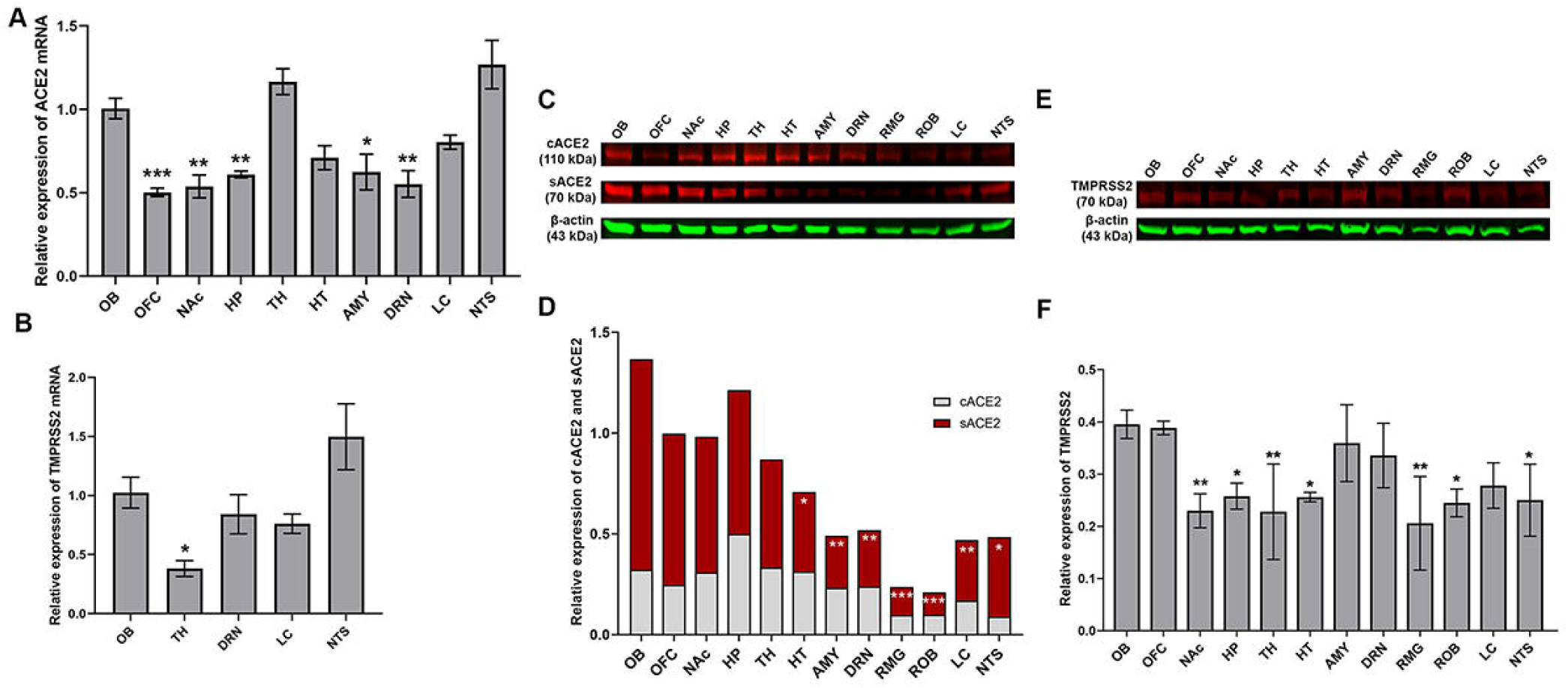
**(A)** Relative mRNA expression of *ACE2* in the orbitofrontal cortex (OFC), nucleus accumbens (NAc), hippocampus (HP), thalamus (TH), hypothalamus (HT), amygdala (AMY), dorsal raphe nucleus (DRN), locus coeruleus (LC), and the nucleus of the solitary tract (NTS) as compared to the olfactory bulb (OB). **(B)** Relative mRNA expression of *TMPRSS2* in OB, TH, DRN, LC, NTS as compared to the OB. **(C and D)** Relative protein expression of cellular ACE2/β-actin and soluble ACE2/β-actin in OB, OFC, NAc, HP, TH, HT, AMY, DRN, raphe magnus (RMG), raphe obscurus (ROB), LC, NTS. **(E and F)** Relative protein expression of TMPRSS2/β-actin in OB, OFC, NAc, HP, TH, HT, AMY, DRN, RMG, ROB, LC, NTS. Values (n = 4 - 5/group) are represented as means (±SEM) and the mRNA data was analyzed by one-way analysis of variance (ANOVA) using Dunnett’s multiple comparisons test (*p < 0.05, **p < 0.01, ***p < 0.001 versus air control). The cACE2 and sACE2 levels were analyzed by two-way ANOVA using Sidak’s multiple comparisons test and the TMPRSS2 protein levels was analyzed by one-way ANOVA using Dunnett’s multiple comparisons test (*p < 0.05, **p < 0.01, ***p < 0.001 versus air control).

### Alcohol exposure impacts ACE2 expression in the mouse olfactory bulb

Our initial results show widespread *ACE2* levels across brain regions, but more limited *TMPRSS2* expression. Next, we turned to immunofluorescent labeling of mouse brain sections to quantify ACE2 and TMPRSS2 expression patterns with finer spatial resolution. Consistent with data from Figure 1, we observe robust ACE2 expression throughout the mouse olfactory bulb (Figure 2A) using fluorescent antibody labeling. ACE2 expression also significantly increased in the olfactory bulb following CIE (Figure 2B). Quantification of immunolabeling shows a significantly increased ACE2 immunoreactive area (t_8_ = 3.084, *p* = 0.0220; Figure 2C). Our data also show expression of TMPRSS2 throughout the olfactory bulb (Figure 2D), but no change in expression following chronic ethanol vapor exposure (t_8_ = 0.2165, *p* = 0.8336; Figure 2E and F).

**Figure 2:**
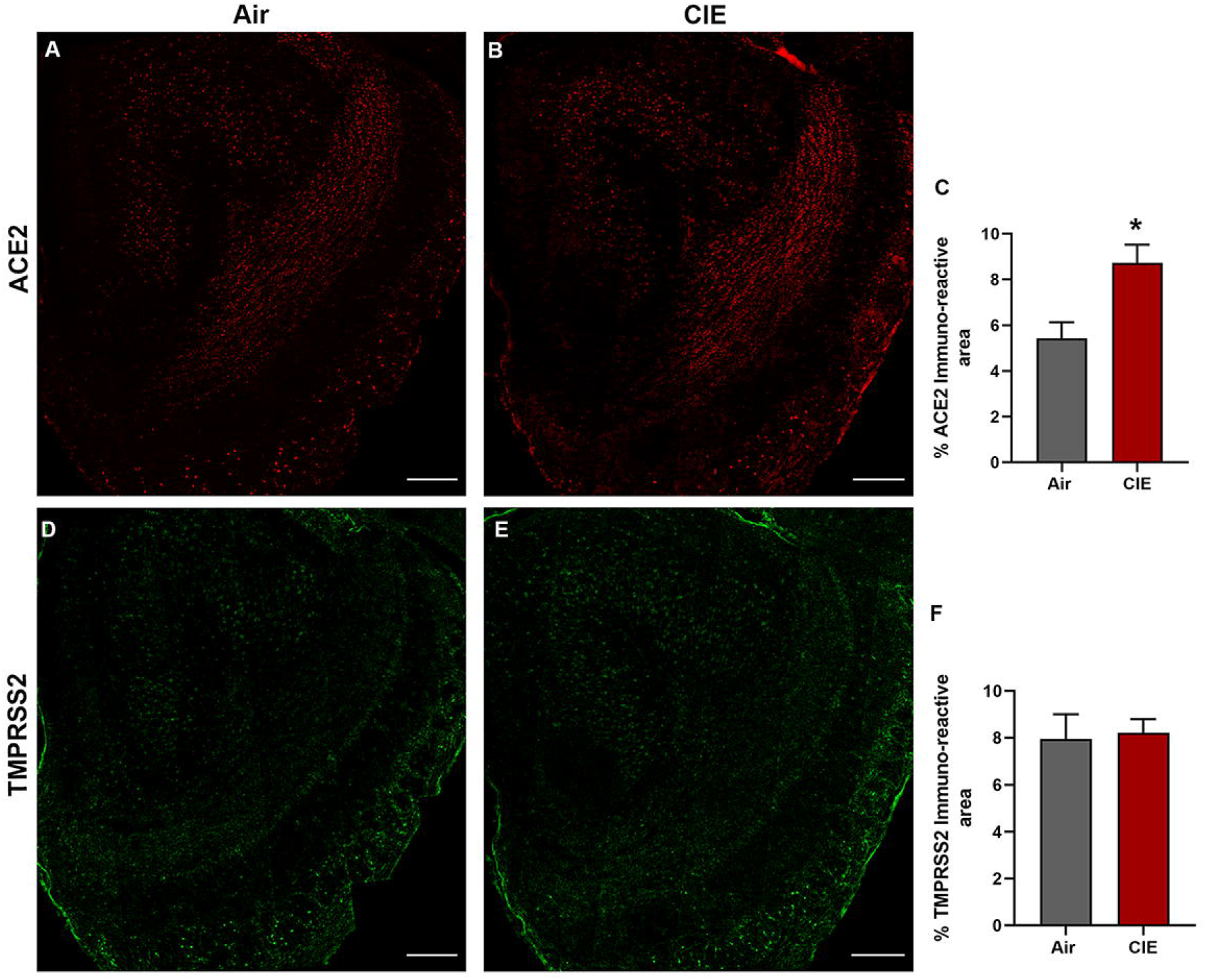
Effect of chronic intermittent ethanol (CIE) exposure on the ACE2 and TMPRSS2 immunoreactivity in the olfactory bulb (OB). Representative confocal images of **(A-B)** ACE2 positive cells in OB of air control and CIE mice. Scale bar = 200 µm **(C)** Graph represents the % ACE2 immunoreactive area in OB. Representative confocal images of **(C-D)** TMPRSS2 in OB of air control and CIE mice. Scale bar = 200 µm **(E)** Graph showing the % TMPRSS2 immunoreactive area in OB. Values (n = 4 - 5/group) are represented as means (±SEM) and the data was analyzed by unpaired student’s t-test with Welch’s correction (*p < 0.05, versus air control).

### Alcohol exposure increased ACE2 levels in discrete hypothalamic regions

The spread of viral expression into several hypothalamic regions has also been reported following viral entry into the olfactory bulb (Netland et al., 2008; Perlman et al., 1990). Hypothalamic projections from olfactory bulb neurons (Nampoothiri et al., 2020; Price et al., 1991; Russo et al., 2018) could mediate this route of viral invasion. With these findings in mind, we tested for differential ACE2 expression levels in hypothalamic regions receiving olfactory bulb projections following CIE. Figure 3 shows robust ACE2 immunolabeling in the paraventricular nucleus of the hypothalamus (PVN), the lateral hypothalamus (LH), or the arcuate nucleus (ARC) (panels A, C, and E, respectively). Consistent with our observations in the olfactory bulb, we observed a significant increase in ACE2 expression in both the PVN (t_9_ = 2.621, *p* = 0.0424) and LH (t_9_ = 3.881, *p* = 0.0100) but observed no changes in the ARC (t_9_ = 1.712, *p* = 0.1449; Figure 3G).

**Figure 3:**
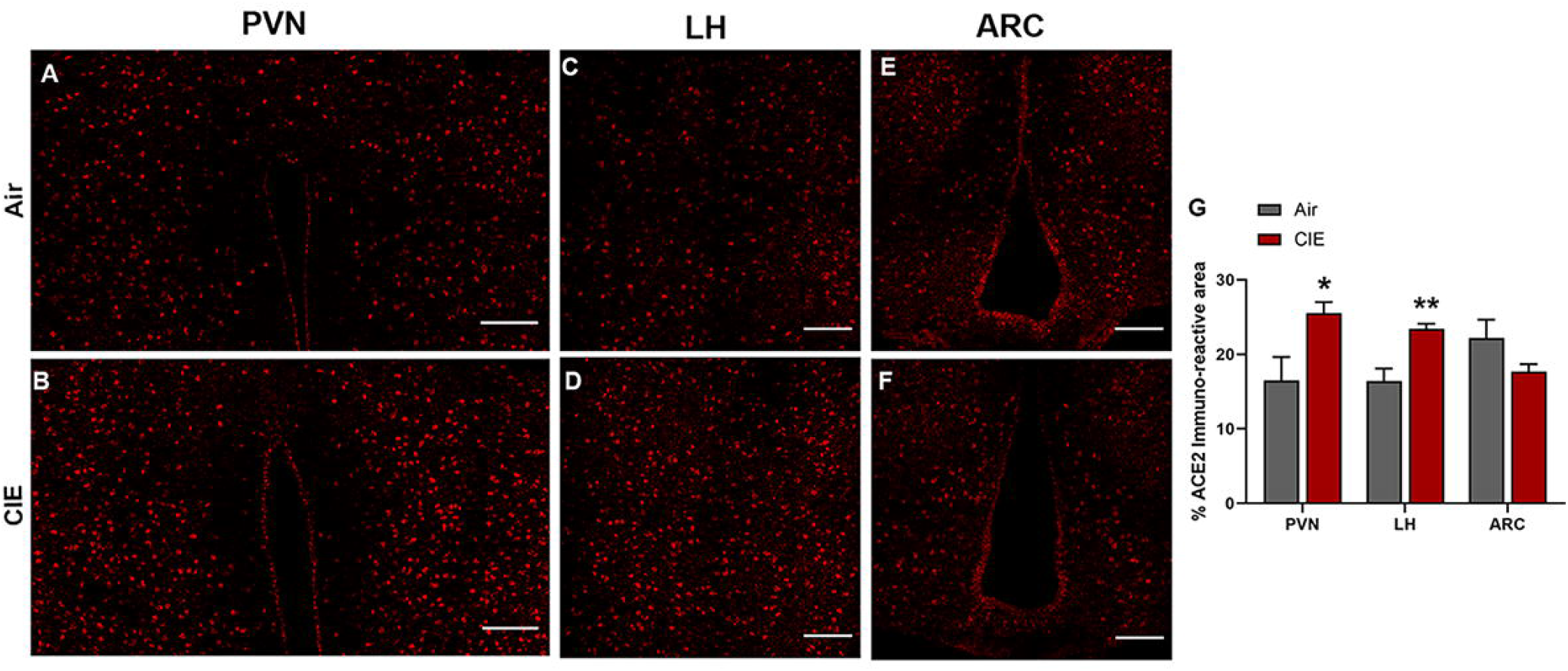
Effect of chronic intermittent ethanol (CIE) exposure on the ACE2 and TMPRSS2 immunoreactivity in different nuclei of hypothalamus–paraventricular nucleus (PVN), lateral hypothalamus (LH), and arcuate nucleus (ARC). Representative confocal images of ACE2 positive cells in **(A-B)** PVN **(C-D)** LH and **(E-F)** ARC of air control and CIE mice. Scale bar = 100 µm **(G)** Graph showing the % ACE2 immunoreactive area in PVN, LH, and ARC. Values (n = 4 - 5/group) are represented as means (±SEM) and the data was analyzed by unpaired student’s t-test with Welch’s correction (*p < 0.05, **p < 0.01 versus air control).

### Differential expression of ACE2 in serotonergic raphe nuclei following CIE exposure

We next utilized our data on CNS-ACE2 expression loci to expand our investigation into regions that could underlie the neurological symptoms associated with SARS-CoV-2. Serotonergic nuclei receive/send projections to the hypothalamus (Papp, 2014) and dysfunction in these nuclei is associated with neurological and affective changes. Therefore, we tested whether dorsal raphe serotonergic neurons are vulnerable to changes in *ACE2* or *TMPRSS2* expression following CIE vapor treatment. We labeled serotonergic neurons by immunostaining for 5-hydroxytryptamine (5HT; Figure 4A) and co-stained for ACE2 (Figure 4B) and TMPRSS2 (Figure 4C). While we observe ACE2 and TMPRSS2 expression throughout the dorsal raphe nucleus, only a subset of serotonergic neurons shows co-labeling for either protein. Further, we observe substantial ACE2 and TMPRSS2 expression in other cell types. In the dorsal raphe, we also observe a significant increase in 5HT immunostaining (t_9_ = 2.865, *p* = 0.0251) and a trend toward increased ACE2 % immunoreactive area (t_9_ = 2.403, *p* = 0.0614), but no significant changes in *TMPRSS2* expression levels (t_9_ = 0.2702, *p* = 0.7964; Figure 5U). Further analysis reveals a significant increase in 5HT/ACE2 colocalization by % immunoreactive area (t_9_ = 2.638, *p* = 0.0354; Figure 5V) and increased Pearson correlation coefficient (t_9_ = 3.494, *p* = 0.0118; Figure 5W). Quantification of Mander’s overlap coefficient (MANDERS et al., 1993) also shows a significant increase in ACE2/5HT colocalization, indicating increased ACE2 expression in 5HT neurons (t_9_ = 3.895, *p* = 0.0117; Figure 5X).

**Figure 4:**
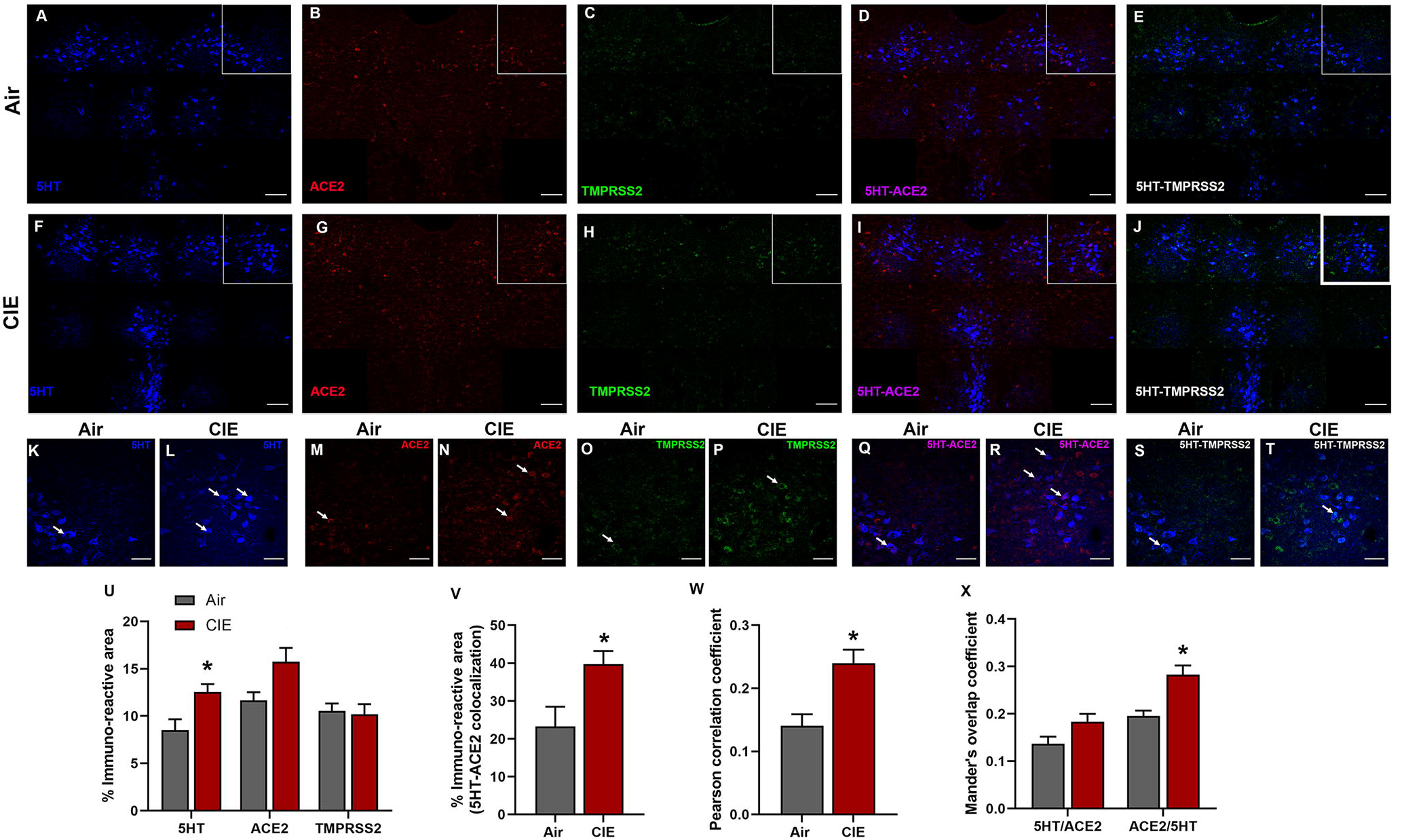
Effect of chronic intermittent ethanol (CIE) exposure on the 5HT, ACE2, and TMPRSS2 immunoreactivity in dorsal raphe nucleus (DRN). Representative confocal images showing **(A and F)** 5HT (blue) **(B and G)** ACE2 (red) **(C and H)** TMPRSS2 (green) **(D and I)** 5HT-ACE2 (magenta) **(E and J)** 5HT-TMPRSS2 (white) positive neurons in DRN of air and CIE mice. Scale bar = 100 µm. Representative single tile confocal images showing **(K and L)** 5HT (blue) **(M and N)** ACE2 (red) **(O and P)** TMPRSS2 (green) **(Q and R)** 5HT-ACE2 (magenta) **(S and T)** 5HT-TMPRSS2 (white) positive neurons in DRN of air control and CIE mice. Scale bar = 50 µm. Graph showing **(U)** % immunoreactive area for 5HT, ACE2, and TMPRSS2, **(V)** % immunoreactive area for 5HT-ACE2 positive neurons, **(W)** Pearson correlation coefficient for 5HT-ACE2 colocalization **(X)** Mander’s overlap coefficient for 5HT-ACE2 colocalization in DRN. Values (n = 4 - 5/group) are represented as means (±SEM) and the data was analyzed by unpaired student’s t-test with Welch’s correction (*p < 0.05 versus air control).

**Figure 5:**
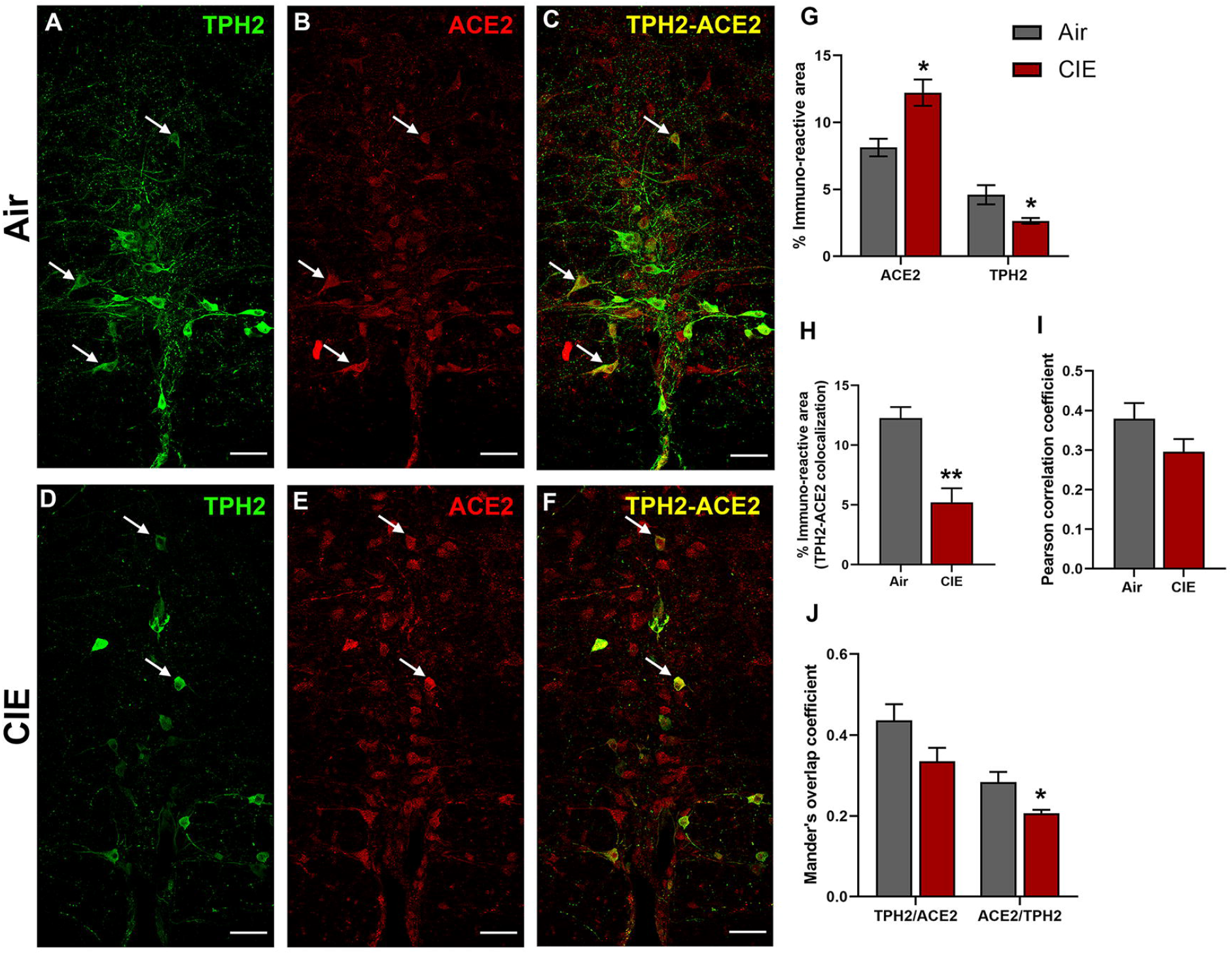
Effect of chronic intermittent ethanol (CIE) exposure on the TPH2 and ACE2 immunoreactivity in raphe magnus (RMG). Representative confocal images showing **(A and D)** TPH2 (green) **(B and E)** ACE2 (red) **(C and F)** TPH2-ACE2 (yellow) positive neurons in RMG of air and CIE mice. Scale bar = 50 µm. Graph showing **(G)** % immunoreactive area for ACE2 and TPH2, **(H)** % immunoreactive area for TPH2-ACE2 positive neurons, **(I)** Pearson correlation coefficient for TPH2-ACE2 colocalization **(J)** Mander’s overlap coefficient for TPH2-ACE2 colocalization in RMG. Values (n = 4 - 5/group) are represented as means (±SEM) and the data was analyzed by unpaired student’s t-test with Welch’s correction (*p < 0.05, **p < 0.01 versus air control).

While the dorsal raphe comprises a majority of CNS 5HT neurons, ACE2 expression in brainstem serotonergic nuclei is highly relevant to SARS-CoV-2 infection as the brainstem is vulnerable to SARS-CoV-2 neuroinvasion (Bulfamante et al., 2021). In this vein, we next tested whether CIE exposure impacts ACE2 expression levels in the raphe magnus. Here we labeled serotonergic neurons using an anti-TPH2 antibody (Figure 5A). We also observed robust ACE2 expression in the raphe magnus, some of which localizes to cells with neuronal morphology (Figure 5B). As in the dorsal raphe, ACE2 is expressed in a subset of raphe magnus serotonergic neurons, with other cell types comprising the remainder of the immunolabeling (Figure 5C). Unlike the dorsal raphe nucleus, ethanol vapor exposure increased ACE2 % immunoreactive area (t_9_ = 3.474, *p* = 0.0153) and decreased TPH2 immunoreactive area (t_9_ = 2.607, *p* = 0.05). Quantification of immunofluorescence also revealed a significant decrease in TPH2/ACE2 co-localization following CIE (t_9_ = 4.730, *p* = 0.0031). Further analysis revealed no change in the Pearson correlation coefficient (t_9_ = 1.647, *p* = 0.1438), but a significant decrease in the ACE2/TPH2 Mander’s overlap coefficient (t_9_ = 2.906, *p* = 0.0349).

Figure 6 shows similar results for the raphe obscurus, another serotonergic brainstem nucleus. Raphe obscurus functions include regulation of breathing (Millhorn, 1986) and gut motility (McCann et al., 1989), both of which are highly relevant to SARS-CoV-2 symptoms. Neuronal projections to the lungs or gut have also been proposed as a route for CNS viral entry (Lima et al., 2020). In Figure 6, we quantified ACE2 expression levels and tested whether CIE impacted localization in the raphe obscurus. We observe robust TPH2 immunolabeling of serotonergic neurons (Figure 6A) and *ACE2* expression (Figure 6B). Only a subset of serotoninergic neurons in the raphe obscurus express *ACE2*, but ACE2 labeling was also widely expressed in non-serotoninergic cells. Indeed, co-labeling of TPH2 and ACE2 was seen in a limited number of neurons (Figure 6C). Based on cellular morphology, other neuronal populations in the raphe obscurus potentially express *ACE2*. Following CIE, TPH2 and ACE2-mediated immunofluorescence increased significantly (Figure 6D and E, respectively), as quantified in Figure 6G (t_9_ = 3.297, *p* = 0.0234 for ACE2 and t_9_ = 4.010, *p* = 0.0051 for TPH2). In addition to an increase in overall levels, we also see increases in metrics of co-expression, including % immunoreactive area (t_9_ = 2.922, *p* = 0.0230; Figure 6H), increased Pearson correlation coefficient (t_8_ = 3.716, *p* = 0.0127; Figure 6I), and both Mander’s overlap coefficients (Figure 6J; t_8_ = 3.190, *p* = 0.0193 and t_8_ = 3.086, *p* = 0.0243 for M1 and M2, respectively).

**Figure 6:**
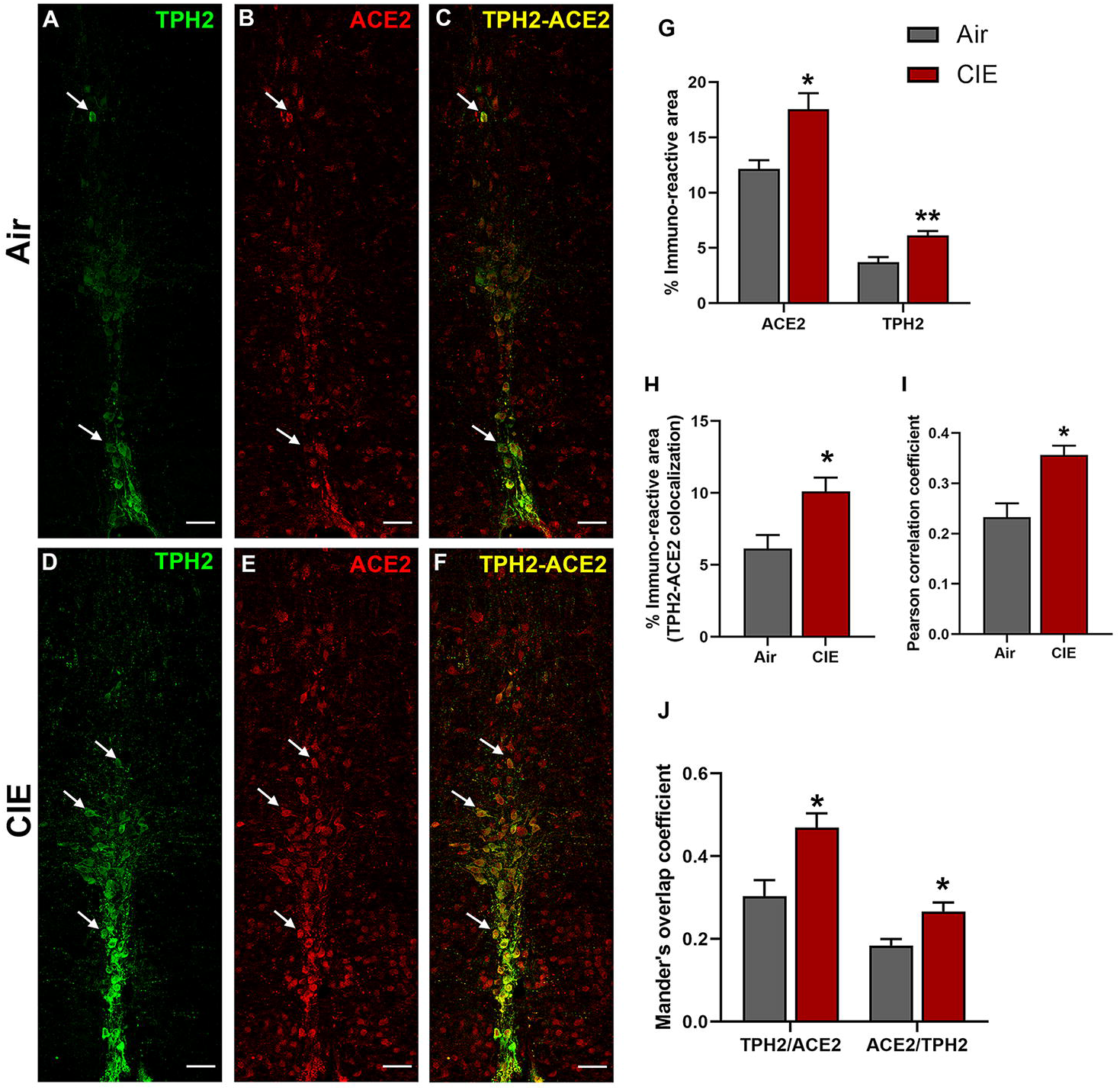
Effect of chronic intermittent ethanol (CIE) exposure on the TPH2 and ACE2 immunoreactivity in raphe obscurus (ROB). Representative confocal images showing **(A and D)** TPH2 (green) **(B and E)** ACE2 (red) **(C and F)** TPH2-ACE2 (yellow) positive neurons in ROB of air control and CIE mice. Scale bar = 50 µm. Graph showing **(G)** % immunoreactive area for ACE2 and TPH2, **(H)** % immunoreactive area for TPH2-ACE2 positive neurons, **(I)** Pearson correlation coefficient for TPH2-ACE2 colocalization **(J)** Mander’s overlap coefficient for TPH2-ACE2 colocalization in ROB. Values (n = 4 - 5/group) are represented as means (±SEM) and the data was analyzed by unpaired student’s t-test with Welch’s correction (*p < 0.05, **p < 0.01 versus air control).

### Impact of alcohol exposure on expression levels in noradrenergic locus coeruleus

After observing differential ACE2 expression in three serotonergic brain regions following CIE, we turned our attention to the locus coeruleus, the primary source of noradrenergic neurons in the CNS that receives projections from the DRN and expresses ACE2 (Figure 1A & D). Because the LC and its noradrenergic projections play significant roles in affective disorders and are impacted by chronic ethanol consumption, we identified noradrenergic neurons by labeling for tyrosine hydroxylase (TH; Figure 7A). Again, we observed ACE2 immunolabeling throughout the LC and in a limited number of TH-positive cells (Figure 7B and C, respectively). Following chronic ethanol exposure, we saw increased labeling of both TH and ACE2 in the locus coeruleus (Figure 7D and E, respectively). Although ACE2 continued to be expressed in TH-positive neurons, there was no indication that this increase in expression was limited to noradrenergic neurons (Figure 7F). Quantification of immunolabeling (Figure 7G) indicates an increased ACE2 expression (t_9_ = 3.504, *p* = 0.0102) and TH expression (t_9_ = 2.964, *p* = 0.0266). Analysis of co-localized area (7H) or Pearson correlation coefficient (7I) did not indicate any changes in co-expression (t_9_ = 1.692, *p* = 0.1377 and t_9_ = 1.533, *p* = 0.1737, respectively). However, Mander’s overlap analysis indicates that a significant increase in ACE2 expression occurs within TH-positive cells, with a trend toward increased TH/ACE2 (t_9_ = 2.279, *p* = 0.0609) and a significant increase in ACE2/TH (t_9_ = 2.475, *p* = 0.0434) expressing cells. Overall, these data show that ACE2 expression in the LC significantly increases following CIE exposure.

**Figure 7:**
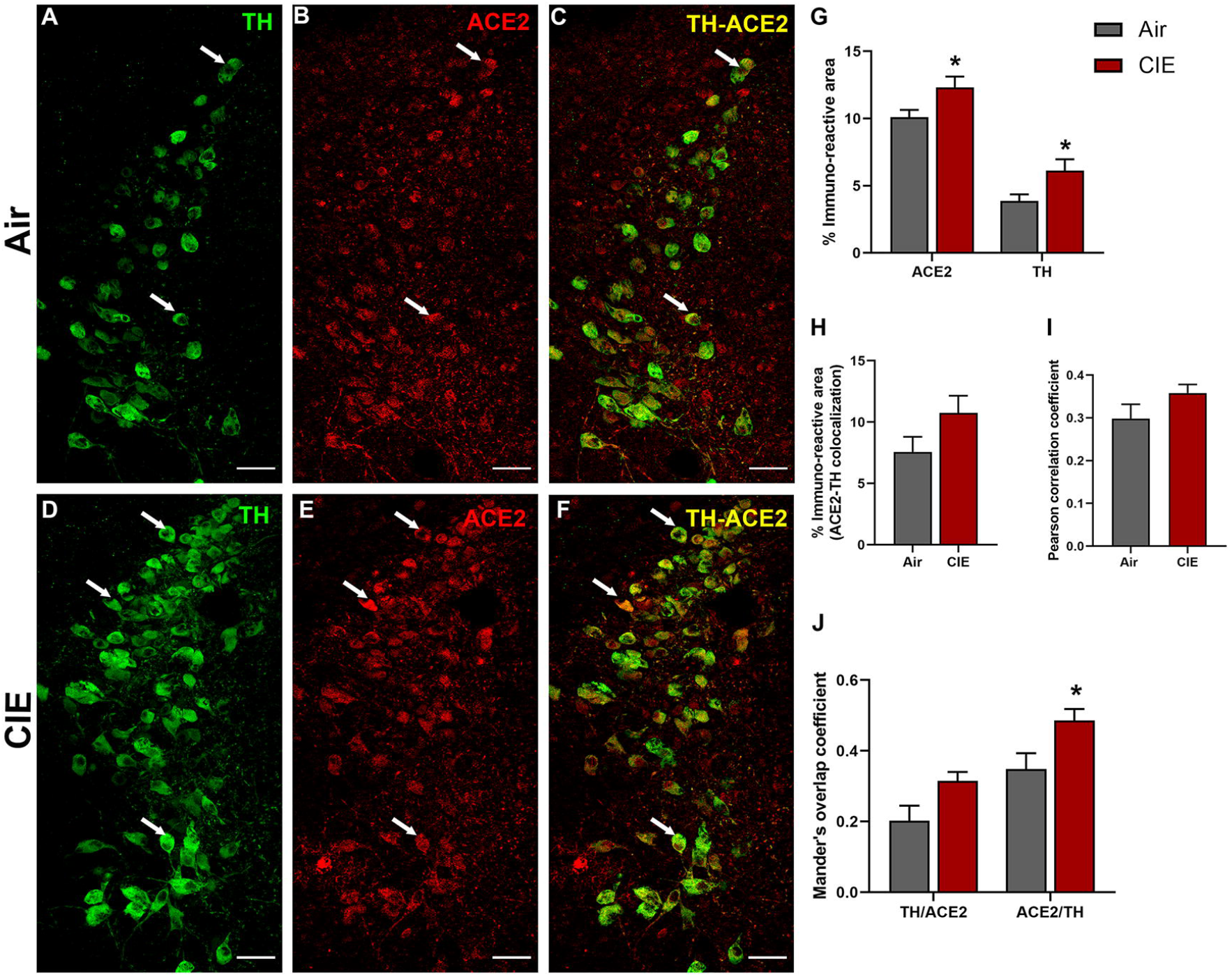
Effect of chronic intermittent ethanol (CIE) exposure on the TH and ACE2 immunoreactivity in locus coeruleus (LC). Representative confocal images showing **(A and D)** TH (green) **(B and E)** ACE2 (red) **(C and F)** TH-ACE2 (yellow) positive neurons in LC of air and CIE mice. Scale bar = 50 µm. Graph showing **(G)** % immunoreactive area for ACE2 and TH, **(H)** % immunoreactive area for TH-ACE2 positive neurons, **(I)** Pearson correlation coefficient for TH-ACE2 colocalization **(J)** Mander’s overlap coefficient for TH-ACE2 colocalization in LC. Values (n = 4 - 5/group) are represented as means (±SEM) and the data was analyzed by unpaired student’s t-test with Welch’s correction (*p < 0.05 versus air control).

### ACE2 levels in the PAG are increased following CIE exposure

The periaqueductal grey (PAG) is vulnerable to SARS-CoV infection (Netland et al., 2008) and impacts critical autonomic functions such as heart rate (Carrive and Bandler, 1991; Lovick, 1992) and breathing (Faull et al., 2019). Our results also showed a significant increase in ACE2 expression in the nearby DRN (Figure 4) following CIE. In investigating this brain region, our initial results showed ACE2 is also expressed in the ventral/lateral (Figure 8A) and dorsal/medial (Figure 8C) PAG. We then tested whether ACE2 levels are also increased in this midbrain locus following CIE. Our results indicate a significant increase in % ACE2 immunoreactive area in the ventral/lateral PAG and a trend toward increased expression in the dorsal/medial PAG (figure 8E; t_8_ = 3.288, *p* = 0.0393 and t_9_ = 2.407, *p* = 0.0577, respectively).

**Figure 8:**
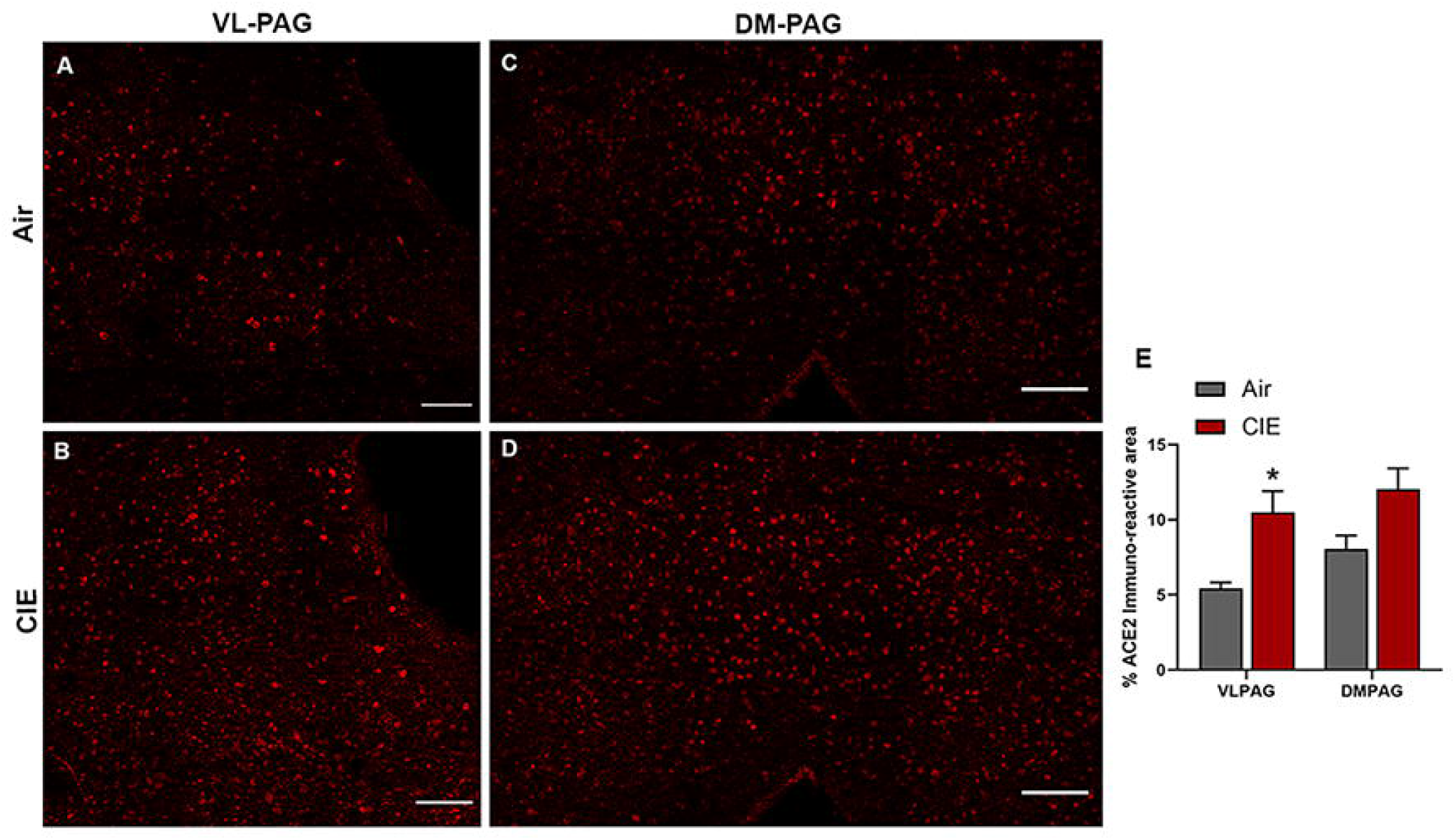
Effect of chronic intermittent ethanol (CIE) exposure on the ACE2 immunoreactivity in the ventrolateral and dorsomedial periaqueductal gray (VL-PAG, DM-PAG). Representative confocal images showing ACE2 positive cells in **(A and B)** VL-PAG **(C and D)** DM-PAG of air and CIE mice. Scale bar = 100 µm. **(E)** Graph showing the % ACE2 immunoreactive area in VLPAG and DMPAG. Values (n = 4 - 5/group) are represented as means (±SEM) and the data was analyzed by unpaired student’s t-test with Welch’s correction (*p < 0.05 versus air control).

### Reduced ACE2 levels in Amygdala following chronic ethanol exposure

Finally, we considered whether alcohol exposure impacts ACE2 expression in the PVT and AMY. Both brain regions receive extensive sensory input and projections from other brain regions. The thalamus is a key node in multiple neuronal circuits and is highly vulnerable to damage from chronic ethanol consumption (Segobin et al., 2019). In both brain regions, we observed robust baseline levels of ACE2 (Figure 9A and C) in control mice. Surprisingly, we saw a trend toward decreased ACE2 expression in the PVT following alcohol exposure (t_9_ = 2.496, *p* = 0.0704) and significantly lower ACE2 levels in the AMY (t_9_ = 2.938, *p* = 0.0364).

**Figure 9:**
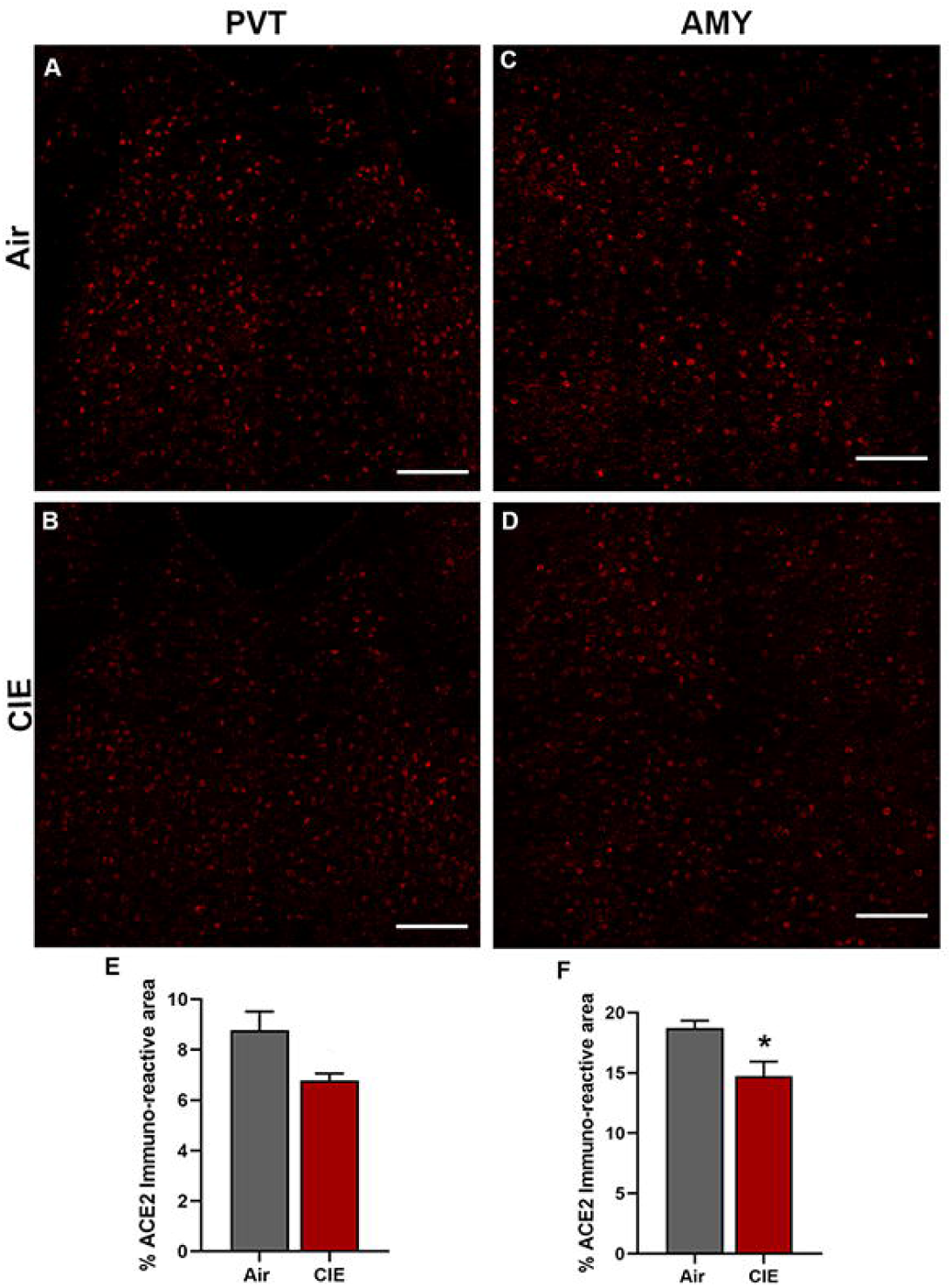
Effect of chronic intermittent ethanol (CIE) exposure on the ACE2 immunoreactivity in the paraventricular thalamic nucleus (PVT) and amygdala (AMY). Representative confocal images showing ACE2 positive cells in **(A and B)** PVT **(C and D)** AMY of air and CIE mice. Scale bar = 100 µm. Graph showing the % ACE2 immunoreactive area in **(E)** PVT **(F)** in AMY. Values (n = 4 - 5/group) are represented as means (±SEM) and the data was analyzed by unpaired student’s t-test with Welch’s correction (*p < 0.05 versus air control).

## Discussion

While SARS-CoV-2 is known to affect the respiratory system, accumulating evidence suggests that it invades the CNS, manifests neurological complications (Siddiqui, 2021), and is even more extensive in AUD and SUD patients (Wang et al., 2021). Moreover, the SARS-CoV-2 associated genes including *ACE2* and *TMPRSS2* were also upregulated in the frontal and motor cortex of chronic alcoholics (Muhammad et al., 2021). Since alcohol abuse increases the risk of respiratory disorder and can drive the risk for COVID-19 infection, in the present study we examined the expression of *ACE2* in mice exposed to 4 weeks of CIE. Our results suggest that ACE2 levels were altered after CIE exposure in various brain regions including the olfactory bulb, hypothalamus, and monoaminergic brainstem regions. While several reports suggest the relevance of addictive disorder to the imbalance in the monoamines in the brain (Jaatinen et al., 2013; Ketcherside et al., 2013; Marcinkiewcz et al., 2016a), the altered ACE2 levels in these neurons may heighten the severity of SARS-CoV-2 infection after chronic alcohol exposure. Elevated ACE2 in the olfactory bulb, hypothalamus, and brainstem might increase the risk of neuroinvasion, while infection of monoaminergic neurons could augment the spread via the widespread network of serotonergic and noradrenergic projections throughout the brain. Altered activity or loss of these neurons may also account for increasing reports of neuropsychiatric complications in the wake of Covid-19 infection.

Based on our results from RT-qPCR and western blot of ACE2, it is apparent that the *ACE2* is expressed throughout the brain, as previously reported (Doobay et al., 2007; Lin et al., 2008; Qiao et al., 2020). ACE2 exists as a transmembrane protein (cACE2) and as a soluble catalytic ectodomain (sACE2), that can be found in plasma and other body fluids including CSF (García-Escobar et al., 2021). A human kidney cell line (HK-2) study showed sACE2-mediated SARS-CoV-2 entry into the cell (Yeung et al., 2021). Interestingly in the present study, we have detected both the isoforms of ACE2 in the brain regions examined and their levels were differentially detected amongst these brain regions. TMPRSS2, a cellular protease that is involved in SARS-CoV-2 virus entry into the cell, is also known to cleave ACE2 thus producing sACE2 (Heurich et al., 2014; Koch et al., 2021). We analyzed TMPRSS2 expression and found that the protein was detected in all the brain regions, but mRNA was below the detection limit in some of the regions examined. Similarly, TMPRSS2 protein was also detected in the human cell line (SH-SY5Y) and mice brain regions including the cortex, hippocampus, striatum, and hypophysis (Qiao et al., 2020). The suggested mechanisms for coronavirus propagation within the brain include internalization in nerve terminals by endocytosis, retrograde transportation, axonal transport, and transsynaptic mode of spreading (Dubé et al., 2018), so high levels of sACE2 in brain regions such as the olfactory bulb may help SARS-CoV-2 propagate into other brain regions.

We next examined the expression levels of ACE2 in the olfactory bulb to assess the potential risk of COVID-19 neuroinvasion after chronic alcohol exposure. COVID-19 patients display anosmia as a primary response to infection and growing evidence of the ACE2 expression in the olfactory bulb confirms the intranasal/olfactory system route is highly relevant to SARS-CoV-2 infection (Burks et al., 2021; Jiao et al., 2021; Qiao et al., 2020; Ueha et al., 2020; Xydakis et al., 2021). An intranasal administration of SARS-CoV-2 in h*ACE2* mice further confirms an infection in sustentacular cells and olfactory neurons (Tang et al., 2021). Interestingly, we observed an increase in the ACE2 and a non-significant change in TMPRSS2 after CIE, suggesting a high risk for SARS-CoV-2 infection in individuals with AUD. Another possible route for viral neuroinvasion could be from the gut through the vagus nerve (Guo et al., 2021; Manosso et al., 2021) and clinical reports show prominent staining for SARS-CoV-2 in the vagus nerve fibers (Bulfamante et al., 2021). Within the CNS, the vagus nerve primarily projects to NTS, releasing excitatory/inhibitory neurotransmitters, acetylcholine, norepinephrine, and neuropeptides. We have observed an abundant expression of ACE2 and TMPRSS2 in NTS as like OB by western blot and RT-qPCR experiments. Further, we have also confirmed the localization of ACE2 in the dorsal motor nucleus of the vagus (DMV) that receives afferent vagal nerve (data not shown). The NTS has widespread efferent pathways to the basal forebrain, amygdala, hippocampus, hypothalamus, and brain stem nuclei (dorsal raphe, raphe magnus, raphe obscurus, and locus coeruleus (Ansari et al., 2007; Henry, 2002). Similar to NTS, olfactory neurons also send/receive projections to other brain nuclei such as the amygdala, hypothalamus, dorsal raphe (Dugué and Mainen, 2009; Murata et al., 2019). Based on the data showing abundant expression of ACE2 in the OB and NTS, which could be a route for SARS-CoV-2 neuroinvasion, we have examined ACE2 levels in other associated regions.

Identification of the SARS-CoV genome in the hypothalamus has also been reported following viral entry into the olfactory bulb (Netland et al., 2008; Pal and Banerjee, 2020; Perlman et al., 1990). Since acute and chronic exposure to alcohol has direct or indirect effects on endocrine functions, which is mediated by the HPA axis (Rachdaoui and Sarkar, 2017), we have examined ACE2 levels in various hypothalamic nuclei and observed that their levels were increased in LH and PVN following CIE. The hypothalamus integrates information from almost all regions of the brain, in particular the brainstem, upon receiving peripheral sensory inputs from different sources including the olfactory system (Nampoothiri et al., 2020; Papp, 2014; Russo et al., 2018).

The majority of serotonergic neurons arise from the DRN, which spans the midbrain and brainstem and plays a significant role in the etiology of neuropsychiatric disorders and AUD (Hasin and Kilcoyne, 2012; López-Figueroa et al., 2004; Marcinkiewcz, 2015; Marcinkiewcz et al., 2016b; Miller et al., 2013; Pivac et al., 2004). Following CIE, we have observed an increase in the 5-HT levels in DR in accordance with a previous study reporting hyperexcitability of DRN neurons after CIE vapor (Lowery-Gionta et al., 2014). Our results also align with a post-mortem study indicating greater TPH immunoreactivity in alcohol-dependent individuals than in control throughout the rostral-caudal axis of DRN (Underwood et al., 2007). The DRN is also known to participate in modulating the core respiratory networks of the brainstem (Bautista et al., 2014; Kaur et al., 2020; Smith et al., 2018), and a recent report indicates the expression of ACE2 in 5-HT neurons in the DRN of rats (Hernández et al., 2021). We examined ACE2 and TMPRSS2 levels in DRN and found that while global ACE2 expression did not change significantly, there was a significant increase in ACE2 colocalization with 5-HT neurons. Although there was no significant change in TMPRSS2 after CIE exposure, the caudal region showed a significant colocalization of 5HT and TMPRSS2.

A clinical study reported immunoreactivity for nucleoprotein (NP) of SARS-CoV-2 in neurons and glial cells of respiratory centers of the brainstem and medulla (Bulfamante et al., 2021). In addition, vagus nerve afferent fibers in the medulla of COVID-19 patients showed intense NP immunostaining, arguing for viral trafficking between the brainstem and the lung or gut. The present study confirms the prominent expression of ACE2 in serotonin and non-serotonin neurons in raphe magnus and obscurus/palladium. It is worth noting that ACE2 expression in these regions was highly conspicuous as compared to the DRN and was significantly increased after CIE exposure. This clinical report and the current preclinical study strongly suggest that ACE2 is prominently expressed in the medullary raphe nucleus and is a viable candidate for SARS-CoV-2 neuroinvasion via the vagal nerve. Interestingly, TPH2 expression was bidirectionally modulated by CIE in these nuclei, with an increase in the TPH2-immunoreactivity in obscurus/palladium and a decrease in the magnus. Because of the difference in the TPH2-immunoreactive area in these regions, the number of TPH2 neurons expressing ACE2 was greater in obscurus/palladium and lesser in magnus in CIE mice. To our knowledge, this is the first report of the effects of chronic alcohol on these medullary raphe nuclei in adult animals. However, in a study of prenatal alcohol exposure, rat pups showed an increase in 5-HT levels in the medullary raphe nucleus at P15 and P21 (Sirieix et al., 2015).

Noradrenergic neurons in the LC have been associated with a wide array of physiological functions including cardiovascular and respiratory control (de Carvalho et al., 2017; Oyamada et al., 1998) via direct projections to the spinal cord or projections to autonomic nuclei like the dorsal motor nucleus (DMV) of the vagus, the rostroventrolateral medulla (RVM), the Edinger-Westphal nucleus, the caudal raphe, the paraventricular nucleus, and the amygdala (Samuels and Szabadi, 2008). As autonomic functions are impaired in COVID-19 patients after infection (Becker, 2021; Milovanovic et al., 2021) and the LC is one of the regions that is associated with these functions, we tested ACE2 immunoreactivity in LC neurons after CIE exposure. Many TH-positive neurons in the LC were positive for ACE2 expression as previously reported (Hernández et al., 2021) and the number of TH-ACE2 neurons was increased in CIE mice, suggesting an increased vulnerability of LC neurons to SARS-CoV-2 infection. The increase in LC-TH expression in the ethanol mice correlates with an increase in norepinephrine levels in urine and CSF samples of individuals during alcohol withdrawal (Hawley et al., 1981; Lubman et al., 1983). The increase in TH expression in LC could be the effect of “kindling” caused by repeated withdrawals of substances as previously reported (Fitzgerald, 2013; LINNOILA, 1987). It is worth noting that there was also abundant ACE2 localization in the Edinger-Westphal nucleus (data not shown), a region involved in the parasympathetic nervous system that receives ascending input from LC (Samuels and Szabadi, 2008).

The PAG is another brainstem region that also plays a critical role in autonomic function, especially breathing (Faull et al., 2019), and shows increased viral antigen for SARS-CoV in h*ACE2* mice after 3 days of intranasal inoculation (Netland et al., 2008). Moreover, acute and chronic alcohol exposure is known to induce robust enhancement of glutamatergic synaptic transmission in dopamine neurons of the VLPAG (Li et al., 2013). We then investigated the effects of CIE on the ACE2 expression levels in VLPAG and DMPAG and found that the ACE2 levels were increased significantly in the VLPAG, suggesting that alcohol can potentiate the vulnerability of this region to viral infection.

The thalamus is a structure of the diencephalon that has extensive nerve connections to the brain stem and the cerebral cortex and is functionally involved in transmitting sensory information to the cortex including taste, which is impaired in COVID-19 patients (Mullol et al., 2020). A few clinical reports have shown a significant role of the thalamus in encephalitis after viral infection (Abel et al., 2020; Beattie et al., 2013; Valencia Sanchez et al., 2021). Moreover, thalamic abnormalities are a key feature in alcohol-related brain dysfunction (Pitel et al., 2015; Segobin et al., 2019). In the present study, we observed decreased ACE2 levels in the PVT following CIE. It is also important to note that the mRNA expression analysis showed a high expression of *ACE2* in the thalamus on par with the OB and NTS. The decreased ACE2 following CIE could be due to shrinkage of the thalamus as previously reported in clinical studies of alcohol dependance (Pitel et al., 2015; Tuladhar and de Leeuw, 2019). Based on the previous clinical reports and the results of the present study, the thalamus is known to have an important role in alcohol dependence and may undergo significant remodeling after SARS-CoV-2 infection.

Besides the brainstem and hypothalamus, ACE2 was also highly expressed in the amygdala (Lukiw et al., 2022; Piras et al., 2021), a brain region implicated in anxiety, stress-related disorders, and the reinforcing effects of alcohol and other drugs of abuse (Roberto et al., 2021). Moreover, RNA-sequencing analysis of amygdala tissue of COVID-19 patients displays upregulation of interferon-related neuroinflammation genes as well as downregulation of synaptic and other neuronal genes (Piras et al., 2021). Based on the COVID-19 reports and the significance of amygdalar circuits in alcohol dependence, we next measured ACE2 levels in the amygdala of CIE mice and found that they were decreased after exposure. Amygdalar ACE2 is known to reduce anxiety by activating Mas receptor signaling (Wang et al., 2016), so this reduction in ACE2 may account for increased anxiety levels in mice after CIE (Ewin et al., 2019).

Altogether, our study provides evidence that chronic alcohol exposure and withdrawal increases *ACE2* expression in monoaminergic neuronal circuits and may potentiate the risk of CNS-viral entry as well as cardiovascular, respiratory, and neuropsychiatric complications. While this study provides a mechanism for increased SARS-CoV-2 risk in individuals who chronically consume alcohol, it represents only an indirect sign of potential vulnerability. Follow-up research is needed to directly show that ethanol-dependent changes in ACE2 levels impact CNS invasion, disease severity, or prevalence of neurological symptoms.

### Disclosure

The authors declare that they have no conflict of interest.

## Acknowledgments

This work was funded through NIH grants R00 AA024215, R00 AA024215-04S2, and R00 AA024215-04S1 to C.M. T.J. was supported by T32 HL007638-35.

