## Supplementary material for "Repeated ethanol exposure and withdrawal alters ACE2 expression in discrete brain regions: Implications for SARS-CoV-2 infection": Tables

**Table 1: List of primers used for quantitating mRNA expression using RT-qPCR**

| **Gene Name** | **Forward/Reverse (5’-3’)** | **Sequence** |
| --- | --- | --- |
| *Ace2* | Forward | TGGGACACGGAGACTTCAGA |
|  | Reverse | TGGCTCCGTTTCTTAGCAGG |
| *Tmprss2* | Forward | TGAAACGCCAGAGCAGGATT |
|  | Reverse | AAATGCCGTCCAGTACCTCG |
| *β-actin* | Forward | CCAGCCTTCCTTCTTGGGTA |
|  | Reverse | GAGGTCTTTACGGATGTCAACG |

**Table 2: Details of the antibodies used in the western blot experiments**

| **Antibodies** | **Use** | **Catalog details** | **Company** | **Dilution** |
| --- | --- | --- | --- | --- |
| ACE2 | Primary | AF933 | Novus Biologicals | 1:250 |
| TMPRSS2 | Primary | sc-515727 | Santa Cruz Biotechnology | 1:1000 |
| β-actin antibody | Primary | sc-47778 | Santa Cruz Biotechnology | 1:2000 |
| IR-dye Anti-Mouse | Secondary | 926-32212 | LI-COR | 1:5000 |
| IR-dye Anti-Goat | Secondary | 926-32212 | LI-COR | 1:5000 |

**Table 3: Details of the antibodies used in immunofluorescence experiments**

| **Antibodies** | **Use** | **Catalog details** | **Company** | **Dilution** |
| --- | --- | --- | --- | --- |
| ACE2 | Primary | AF933 | Novus Biologicals | 1:250 |
| TMPRSS2 | Primary | sc-515727 | Santacruz | 1:300 |
| TPH2 | Primary | NB100-7455 | Novus Biologicals | 1:1000 |
| 5HT | Primary | 20080 | Immunostar | 1:4000 |
| TH | Primary | MAB-318 | Millipore | 1:200 |
| Donkey Anti-Goat | Secondary | 705-165-147 | Jackson Immunoresearch laboratories | 1:800 |
| Donkey Anti-Mouse | Secondary | 715-545-150 | Jackson Immunoresearch laboratories | 1:800 |
| Donkey Anti-Rabbit | Secondary | A31573 | Invitrogen | 1:800 |
| Donkey Anti-Rabbit | Secondary | 711-475-152 | Jackson Immunoresearch laboratories | 1:800 |
